# The nuclear structural protein NuMA is required for efficient DNA replication by ensuring association of core replication factors to replication forks

**DOI:** 10.64898/2026.05.09.723787

**Authors:** Zhen-Guo Wang, Sara Knowles, Fang Wang, Weihang Chai

## Abstract

DNA replication is a strictly regulated process during cell proliferation to ensure faithful duplication of the genome. Its firing and elongation can be arrested or temporally inhibited in response to a variety of internal and external causes. Inside cells numerous factors including cell cycle checkpoints, protein kinases, and others are involved in the control of this process to maintain genome integrity. Here, we describe that NuMA, a nuclear scaffolding protein, plays an important role in regulating DNA replication. We show that NuMA is present at active replication forks, and its deficiency impairs cell viability, reduces the replication fork speed and increases origin firing, leading to increased level of γH2AX and the activation of the ATM-CHK2 DNA damage response pathway. Mechanistically, our results show that NuMA depletion reduced the association of multiple key replisome proteins to replication forks, suggesting that NuMA is essential for efficient replisome proteins binding to ongoing forks. Our study uncovers a novel function of NuMA in maintaining genome stability, providing new insights into the important role of nuclear structural proteins in safeguarding DNA replication.

## Introduction

DNA replication is a highly regulated, multi-step process essential for accurate genome duplication during each cell cycle to ensure faithful transmission of genetic information during cellular proliferation. It requires the coordinated action of multiple proteins and regulatory factors. Dysfunction of these proteins destabilizes DNA replication and promotes genome instability, contributing to a number of developmental disorders and cancer (Nordman and Orr-Weaver 2012, Jackson, Laskey et al. 2014, Munoz and Mendez 2017).

Eukaryotic DNA replication depends on the precise assembly and coordination of replisome activities at replication forks. Once the S phase begins, the inactive MCM2-7 helicase complexes that were previously loaded at licensed origins are activated and the active replicative helicase CMG (Cdc45/MCM2-7/GINS) is assembled. CMG unwinds the parental DNA duplex ahead of the fork, producing single-stranded (ss) DNA that is rapidly coated by RPA to protect it from nucleolytic degradation and to facilitate the recruitment of additional fork-associated factors. DNA synthesis is then carried out by DNA polymerase α (POLα) and POLδ for lagging-strand synthesis, while POLε is primarily responsible for leading-strand synthesis. PCNA, the sliding clamp, is loaded by RFC and confers high processivity to both polymerases. Numerous replisome components, including AND-1, MCM10, CLASPIN, and DNA topoisomerases, coordinate the coupling between helicases and polymerases, maintain fork stability, and ensure proper resolving of super helical tension ahead of the moving fork (Costa and Diffley 2022).

Studies have shown that potential origins are much more abundant than those activated (fired) during each S phase. The dormant origins remain unfired under normal conditions but serve as a critical backup system. When forks slow or stall due to replication stress, these dormant origins can be activated locally to ensure timely duplication of the affected regions and prevent the formation of under-replicated DNA. The availability and chromatin distribution of dormant origins therefore represent an essential layer of support that maintains the robustness of replication under both physiological and stress conditions. However, the mechanisms controlling DNA replication speed and the choice and efficiency of replication origins remain poorly understood (Gilbert 2010, Mechali 2010).

Replication fork progression is constantly monitored by the ATM and ATR DNA damage response checkpoint pathways (Shechter, Costanzo et al. 2004). ATR is activated by stretches of RPA-coated ssDNA generated at perturbed forks and signals through CHK1 to stabilize the replisome. Activation of ATR prevents inappropriate processing of stalled forks, and suppresses unscheduled remodeling events that would threaten fork integrity. ATR does not participate in the firing of replication origins, but limits unnecessary activation of dormant replication origins and prevents the accumulation of DNA damage during the S phase (Saldivar, Cortez et al. 2017). ATM responds to double-strand breaks (DSBs) that may arise from fork collapse and signals through CHK2 to modulate CDK activity and maintain controlled S-phase progression (Falck, Mailand et al. 2001, Sorensen, Syljuasen et al. 2003, Kang, Wei et al. 2008, Honaker and Piwnica-Worms 2010). These responses collectively preserve fork stability, promote fork restart, and prevent replication intermediates from converting into deleterious DNA lesions.

Emerging evidence suggests that nuclear organization plays a critical role in regulating DNA replication by compartmentalizing the genome into early- and late-replicating zones, thereby controlling the timing of origin firing and fork progression (Jackson 2005, Muck and Zink 2009, Van Bortle and Corces 2012, Marks, Smith et al. 2016). Euchromatin located in the nuclear interior generally replicates early in the S phase, whereas condensed heterochromatin at the periphery replicates late. This spatial organization ensures orderly and efficient genome duplication while preserving genome stability. The nuclear lamina, composed of meshwork of proteins (lamins) underlying the nuclear envelop, serves as a structural scaffold that anchors replication foci and chromatin. Disruption of lamin organization has been shown to impair DNA synthesis progression (Moir, Spann et al. 2000). In addition, RIF1 (Rap1-interacting factor 1) functions as a molecular hub that coordinates 3D genome organization with replication timing, helping to establish boundaries between early- and late-replicating domains through modulation of chromatin looping (Smith and Aladjem 2014, Gnan, Flyamer et al. 2021). Cohesin, together with CTCF, further contributes to genome organization by anchoring DNA to the nuclear matrix and creating loops that control active origin spacing (Li, Huang et al. 2013). Despite these advances, the mechanisms by which nuclear structural components regulate DNA replication remains largely unknown.

The nuclear mitotic apparatus (NuMA) is a structural component of nuclear matrix. It is a high molecular weight (238 kDa) protein (2115 amino acids) comprised of N-terminal and C-terminal globular domains and a central long coiled-coil domain (Yang, Lambie et al. 1992). While NuMA is best known for its crucial role in spindle assembly and organization during mitosis (Radulescu and Cleveland 2010, Kiyomitsu and Boerner 2021), it is also localized in the interphase nucleus and associates with chromatin via the evolutionarily conserved sequence in its C-terminus (Abad, Lewis et al. 2007, Rajeevan, Keshri et al. 2020, Serra-Marques, Houtekamer et al. 2020). Emerging evidence has revealed the critical role of NuMA in the nucleus that regulates transcription, DNA repair and chromatin organization. NuMA promotes the mechanical robustness of the nucleus (Serra-Marques, Houtekamer et al. 2020) and affects gene expression (Harborth, Weber et al. 2000, Ohata, Miyazaki et al. 2013). In addition, it has been proposed that NuMA may promote homologous recombination repair by regulating the accumulation of the ISWI ATPase SNF2h at DNA breaks and serving as a negative regulator of 53BP1 (Vidi, Liu et al. 2014, Salvador Moreno, Liu et al. 2019). Recently, NuMA has been found to bind to RNA polymerase II and TDP1 at gene promoters to protect cells from oxidative damage (Ray, Abugable et al. 2022), and ADP-ribosylation of NuMA promotes single-strand break repair and transcription following oxidative stress (Abugable, Liao et al. 2025). Moreover, NuMA regulates ultraviolet (UV)-induced DNA lesion repair, and depletion of NuMA resulted in delayed UV lesion repair and increased cellular UV sensitivity (Kim, Kim et al. 2025). Despite emerging evidence implicating NuMA in DNA repair, its role in genome maintenance during DNA replication remains unexplored.

In this study, we report that NuMA localizes at active DNA replication forks under unperturbed condition, and its depletion significantly impairs replication progression, causing reduced replication speed and increased origin firing. NuMA depletion results in elevated γH2AX levels, micronuclei formation, and the activation of the ATM/CHK2 DNA damage response pathway, indicating that the impaired DNA replication increased DNA damage due to DSB formation. Consequently, cell viability is substantially impaired by NuMA loss. Consistent with prior findings, NuMA depletion results in pronounced nuclear deformation, further supporting its role in preserving the nuclear architecture. Furthermore, we find that NuMA is important for promoting the association of various DNA replication factors including PCNA, DNA polymerases and MCMs to replication forks. Together, these findings uncover a novel role of NuMA in regulating DNA replication and cell viability, highlighting its essential function in sustaining replication fork integrity and genomic stability.

## RESULTS

### NuMA loss impairs cell viability

To investigate the potential role of NuMA in regulating DNA replication, we first analyzed whether loss of NuMA affected cell viability. We used the published HeLa cell lines in which NuMA can be conditionally depleted (referred to as CTT20, H11.2 and H12.2 in this study) (McKinley and Cheeseman 2017). These cells contain a doxycycline (Dox)-inducible Cas9 gene incorporated into the genome. The CTT20 cells express control sgRNA, whereas H11.2 and H12.2 cells express individual sgRNA targeting distinct sites in early exons of NuMA. Upon Cas9 induction by Dox treatment, the NuMA gene can be knocked out in H11.2 and H12.2 cells. Using these cells, we observed significant reduction of NuMA proteins following 4 days of Dox treatment (Fig. 1A). However, we did not observe complete elimination of NuMA protein, probably because some cells might have escaped the Cas9 cutting or the NuMA gene break was repaired in the way that retained the open reading frame. Extending Dox treatment beyond 4 days did not further reduce NuMA protein levels (data not shown). We assessed cell viability using the colony formation assay, and observed that while doxycycline treatment modestly reduced the viability control cell CTT20, NuMA depletion substantially reduced the viability of H11.2 and H12.2 (Fig. 1B, Suppl Fig. S1).

**Fig. 1.**
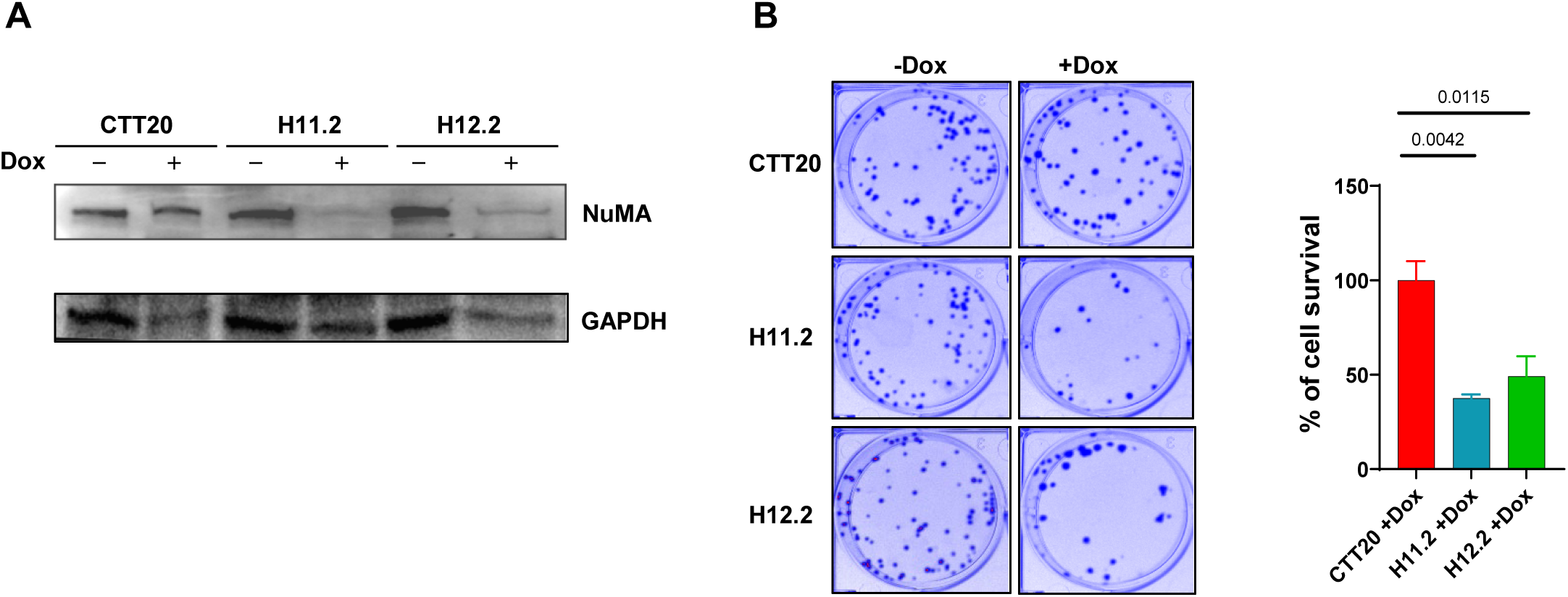
NuMA deficiency decreases cell viability. (A) Western blot showing NuMA depletion after Dox induction of Cas9 in CTT20, H11.2 and H12.2 cells. (B) Colony formation images and bar graph depicting the average percentage of cell numbers relative of no-Dox treated control. Data without normalization is shown in Suppl Fig. S1. Representative images were shown from at least two independent experiments. Ordinary one-way ANOVA was performed for analysis. Error bars represent standard error of the mean (SEM) from three replicates.

### NuMA localizes at replication forks under normal conditions

NuMA, besides its crucial role in spindle assembly and organization during mitosis, is a structural component of nuclear matrix and also associates with chromatin (Abad, Lewis et al. 2007, Rajeevan, Keshri et al. 2020, Serra-Marques, Houtekamer et al. 2020). Immunostaining showed that NuMA predominantly localized in the nucleus (Fig. 2A). We then use the iPOND (isolation of proteins on nascent DNA) assay (Dungrawala and Cortez 2015) to determine whether NuMA associates with replicating DNA (Fig. 2B). In iPOND, cells were pulse-labeled with thymidine analog 5-ethynyl-2’-deoxyuridine (EdU) for 10 min to allow EdU incorporation into the newly synthesized DNA. Following formaldehyde crosslinking of DNA with associated proteins, biotin was attached to EdU via the click chemistry, and the biotinylated nascent DNA was then captured by streptavidin beads from lysed and sonicated cells. Proteins that were physically close to replication forks at the time of EdU labeling were co-purified and then identified by Western blotting. We detected NuMA in the beads fraction in the control CTT20 cells (Fig. 2C). The NuMA iPOND signal was substantially reduced after NuMA depletion, as expected, suggesting that the iPOND signal was specific to NuMA (Fig. 2C).

**Fig. 2.**
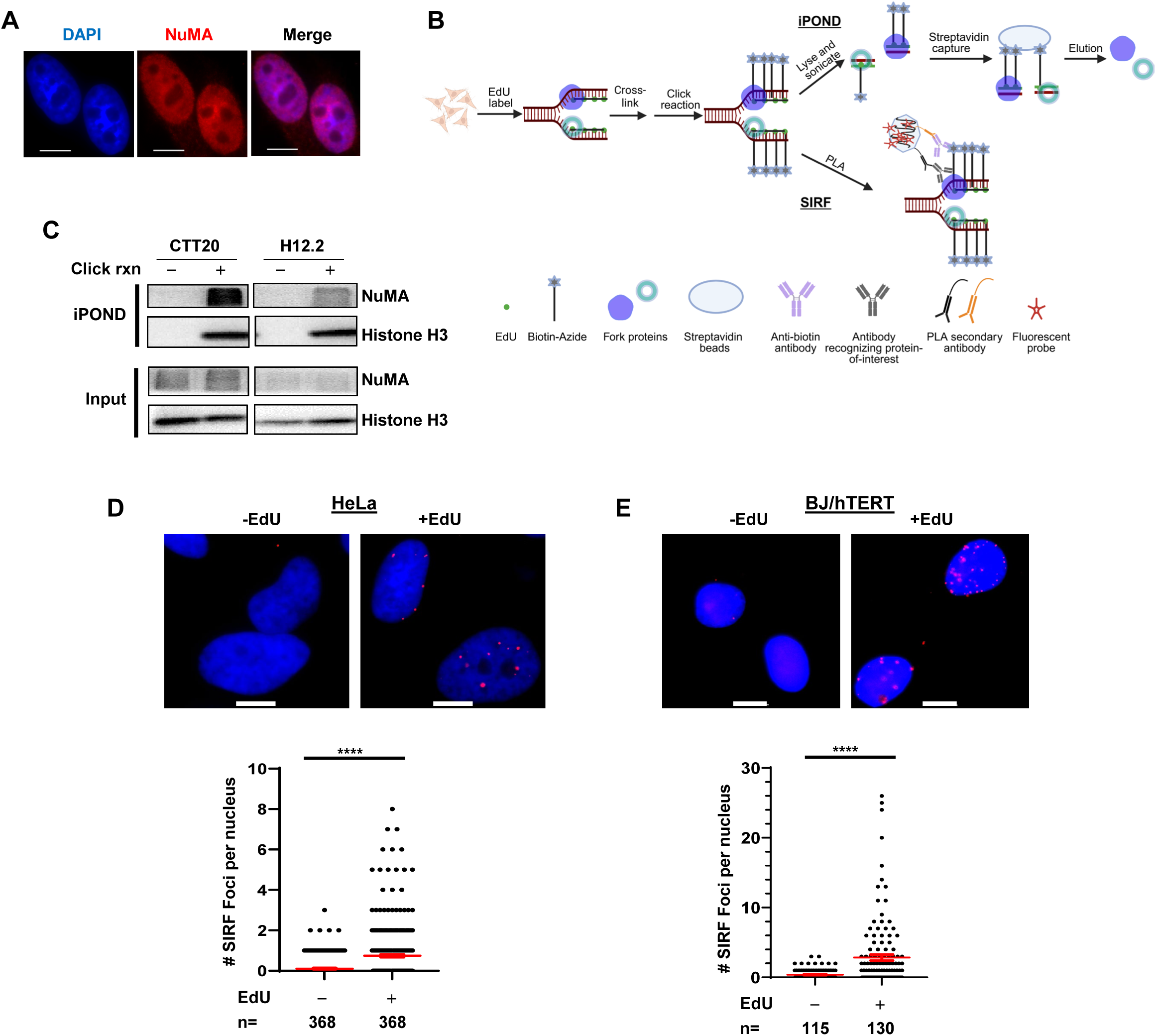
NuMA localizes to replication forks and affects DNA replication. (A) IF shows NuMA localization in the nucleus. HeLa cells were stained with anti-NuMA antibody and counterstained with DAPI. Scale bars: 10 µm. (B) Scheme of iPOND and SIRF assay. Nascent DNA is pulse-labeled with EdU, followed by crosslinking (iPOND) or fixation (SIRF). The click reaction is performed to convert EdU to a detectable biotin conjugate. In the iPOND assay, cells are lysed and chromatin is sonicated, followed by streptavidin beads incubation to capture biotinylated DNA and its associated proteins. Proteins are then eluted from beads and detected by Western blotting. In the SIRF assay, cells are incubated with anti-biotin and anti-target protein antibodies, followed by PLA amplification to visualize the localization of the target protein in close proximity with EdU-labeled forks. (C) iPOND assay showing NuMA localization at active replication forks in CTT20 cells. Cells were grown in the presence of Dox (1 µg/ml) for 4 days without HU treatment, crosslinked, and collected for iPOND analysis. No click reaction was the negative control. Histone H3 was the positive control. The experiments were done at least two times and representative blots were shown. NuMA deletion in H12.2 cells dramatically decreased NuMA iPOND signaling, supporting that the NuMA iPOND signal was specific. (D) Detection of NuMA at active forks with SIRF in HeLa and BJ/hTERT cells. Cells were labeled with EdU for 8 min and SIRF was performed. Representative SIRF images quantification of NuMA signals are shown. Scale bars: 10 µm. The number of SIRF foci per nucleus were manually counted. Two independent experiments were performed and scatter plots from one experiment are shown in the figure. n: the number of cells analyzed in each sample. *P* values were calculated using Mann-Whitney test. **** *P*< 0.0001.

To ensure that the observed NuMA association with replication forks was not specific to the CTT20 cell line, we then checked NuMA association with forks using two other cells lines: HeLa (ATCC# CCL-2, referred to as HeLa in this study) and the normal skin fibroblast BJ immortalized with telomerase (BJ/hTERT). We performed SIRF (*in situ* protein interactions at nascent and stalled replication forks) assays (Roy, Luzwick et al. 2018), which offers sensitive visualization of protein localization at forks at a single-cell level if the protein-of-interest is in close proximity (<40 nm) to EdU-labeled nascent strands. The SIRF assay is similar to iPOND assay in the initial EdU labeling and biotin conjugation steps, but during later steps, signals were amplified inside single cells through Proximity Ligation Assay (PLA) to visualize the colocalization of the target protein with biotinylated EdU (Fig. 2B). Very few SIRF positive cells were detected when EdU was omitted (Figs. 2D, 2E), suggesting that SIRF assay was specific for determining protein localization at forks. With EdU incorporation, NuMA SIRF foci increased in both HeLa and BJ/hTERT cells, indicating its presence at or near replication forks in both cancer and non-cancer cell lines (Figs. 2D, 2E).

### NuMA deficiency impairs DNA replication progression under normal conditions

Since NuMA could be found at replication forks, we then determined whether NuMA played a role in regulating DNA replication using the DNA fiber assay. NuMA was depleted with two individual siRNAs for 48 hrs (Fig. 3A) and replication tracks were then sequentially pulse-labeled with the thymidine analogues chlorodeoxyuridine (CldU) and iododeoxyuridine (IdU) for 20 min each. Cells were lysed and DNA fibers were stretched onto glass slides and used for immunostaining with specific antibodies against CldU and IdU (Fig. 3B). We observed that DNA fiber lengths were shortened for both CldU and IdU fiber tracks after NuMA knockdown (Figs. 3B, 3C). The average replication fork speed was ∼1.2 kb/min in control cells, and 0.9 kb/min and 0.7 kb/min in NuMA knockdown cells after targeting NuMA with two different siRNA (siNuMA-1 and siNuMA-2), indicating that loss of NuMA impaired replication progression.

**Fig. 3.**
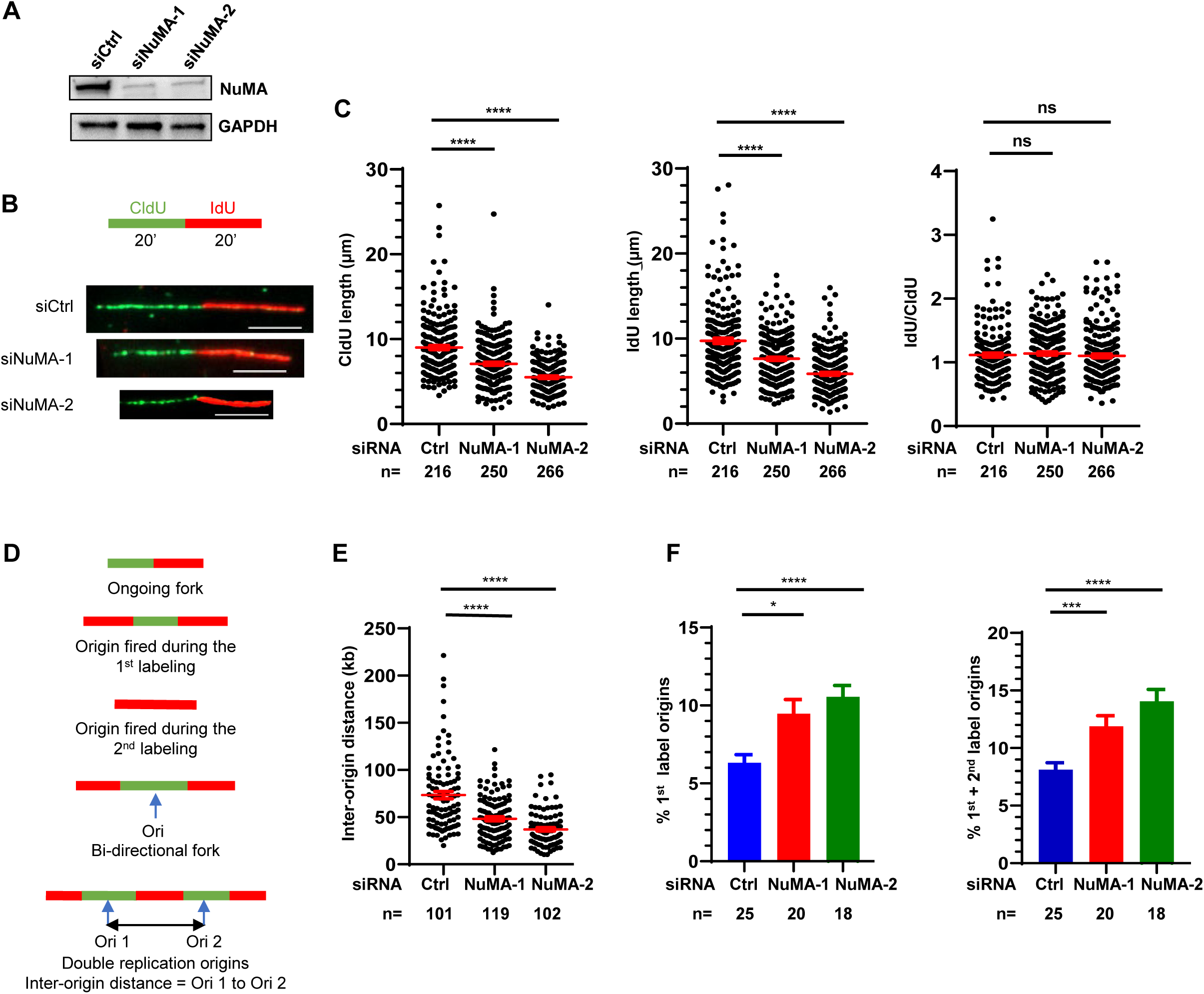
NuMA is critical for maintaining DNA replication progression and controlling origin firing. (A) Western blot showing the knockdown of NuMA by two different siRNAs in HeLa. (B) Scheme of DNA fiber analysis and representative images of DNA fiber. Scale bars: 5µm. (C) DNA fiber analysis of control (Ctrl) and NuMA knockdown HeLa cells under normal replication conditions. CldU and IdU lengths and the ratio of IdU/CldU were calculated for each condition. To obtain fork speed in kb/min, the length of CldU or IdU tracks in kb was divided by the labeling time (20’). Fiber length values were converted into kb using the conversion factor 1 µm = 2.59 kb (Jackson and Pombo 1998). n: the number of fibers counted. *P* values were calculated using one-way ANOVA. **** *P*< 0.0001; *** *P*<0.001; * *P*<0.05; ns (not significant) *P*>0.05. Red lines indicate the mean values. Error bars: SEM. Two independent experiments were performed and scatter plots from one experiment are shown. (D) Schematic representations of DNA fiber patterns showing the 1st label origins, the 2nd label origins, bi-directional fork, and inter-origin distance. (E) Inter-origin distance measurement upon NuMA depletion. *P* values were calculated using one-way ANOVA. **** *P* < 0.0001. n: the number of IODs counted. Red lines indicate the mean values. Error bars: SEM. Two independent experiments were performed and scatter plots from one experiment are shown in the figure. (F) Percent of 1st label origins and combined the 1st and 2nd label origins in NuMA knockdown cells. n: the number of each individual microscopic image, which is comprised of around 50 green tracks. Error bars: SEM. *P* values were calculated using one-way ANOVA.

It has been observed that fork rate is inversely correlated to the number of active forks, perhaps due to constrains on a limiting factor required for replication, such as the dNTP pool (Herrick and Sclavi 2007, Herrick and Bensimon 2008, Poli, Tsaponina et al. 2012). The reduced fork speed in NuMA knockdown cells prompted us to determine whether origin firing was increased after NuMA knockdown. We analyzed the inter-origin distance (IOD) by measuring the distance between the midpoints of the two neighboring origins (green tracks) from Red-Green-Red-Green-Red fibers (Fig. 3D). The average IOD in control cells was 73 kb, within the reported range from 40 kb to 144 kb due to clonal variation (Jackson and Pombo 1998, Guilbaud, Rappailles et al. 2011). In contrast, the siNuMA-1 and siNuMA-2 knockdown cells showed an average IOD of 48 kb and 37 kb, respectively (Fig. 3E). This markedly shortened IOD suggested an increased dormant origin firing after NuMA depletion. We also calculated the percentage of origins during the first pulse labeling and the combined first and second pulses of labeling. Considering that there may be ambiguous pattern of fibers, we only considered red-green-red as one active 1st origin for simplicity. The red-only fiber was considered as the 2nd labeling origin. NuMA-deficient cells showed increased percentages of both the 1st label origins and the combined 1st and 2nd label origins compared to control cells, further supporting an excessive firing of dormant origins after NuMA knockdown (Fig. 3F).

### NuMA deficiency induces DNA damage and causes nuclear deformation

Histone protein H2AX phosphorylation at S139 (termed γH2AX) is a marker for DNA damage, particularly DSBs (Kinner, Wu et al. 2008). DSBs are normally visualized by the formation of γH2AX foci as these two events show close (1:1) correlation (Redon, Dickey et al. 2009). We observed a significant increase of γH2AX level after Cas9 induction to remove NuMA in H12.2 cells (Fig. 4A). Elevated γH2AX level was also detected in HeLa cells after NuMA knockdown using two different siRNAs (Fig. 4B). Consistently, immunostaining showed a markedly increased number of γH2AX foci in H12.2 cells upon NuMA deletion (Fig. 4C). Since γH2AX can accumulate during normal S phase in the absence of exogenous DNA-damaging agents as well as in response to replication stress, we examined whether NuMA depletion increased the S phase of the cell cycle. Cell cycle analysis showed that NuMA depletion slightly reduced the S phase population (Fig. 4D, Suppl Fig. S2). These observations suggest that γH2AX accumulation observed in NuMA deficient cells was not due to S phase cell increase. In addition, we observed a slightly increased phosphorylation of CHK2 (p-Thr68) after NuMA depletion, implying the activation of the ATM-CHK2 pathway in response to DSB formation (Fig. 4A). Such ATM/CHK2 activation was unlikely due to fork collapse, as no increase of fork collapse was observed in NuMA deficient cells (Suppl Fig. S3).

**Fig. 4.**
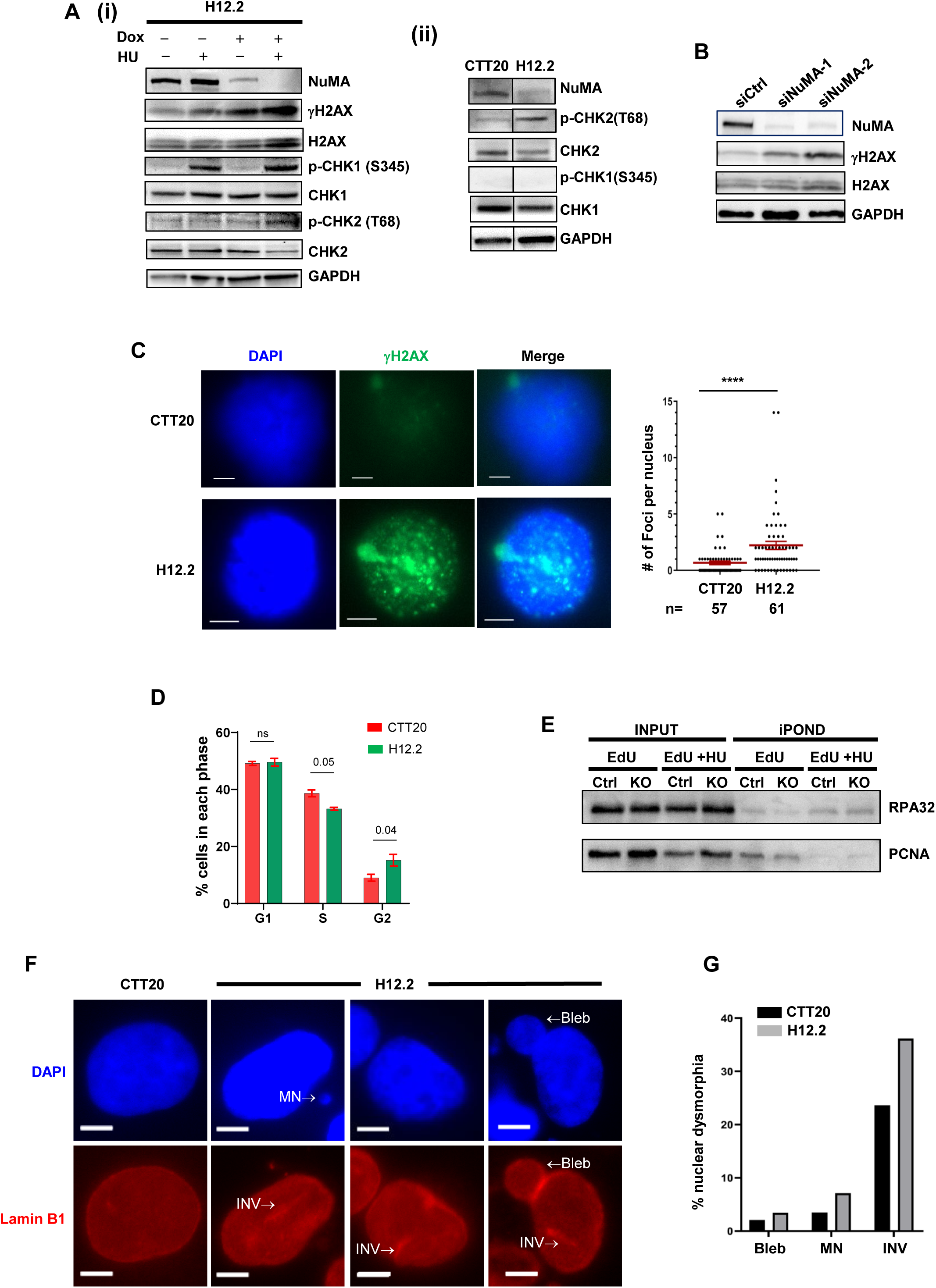
NuMA deficiency increases DNA damage. (A) Western blot showing increased γH2AX and p-CHK2 levels but not p-CHK1 in NuMA-deficient cells. (i) CTT20 and H12.2 cells were grown ± Dox (1 µg/ml) for 4 days and treated with or without HU (4 mM) for 3 hours. (ii) CTT20 and H12.2 cells were treated with Dox (1 µg/ml) for 4 days and collected for western blot analysis. Experiments were repeated at least three times and representative blots are shown. (B) Increased γH2AX in HeLa cells transfected with NuMA siRNAs. HeLa cells were transfected with two different siRNAs for NuMA or control siRNA. Three days later, cells were trypsinized and used for Western blot analysis. (C) IF showing the increased γH2AX foci in NuMA-depleted H12.2 cells. Cells were stained with antibody against γH2AX and counterstained with DAPI. The number of γH2AX foci per nuclei was manually counted. Scale bars: 5 µm. n: the number of nuclei counted. Error bars: SEM. *P* values: Mann-Whitney test. **** *P*< 0.0001. (D) Percentage of cells in each cell cycle phase after NuMA depletion. CTT20 and H12.2 cells were grown in the presence of Dox (1 µg/ml) for 4 days and were collected for cell cycle analysis using flow cytometry. The averages of percentage of cells in each phase were calculated from two experiments. Error bars: SEM. *P* values: multiple paired *t* tests. (E) iPOND analysis of RPA32 and PCNA. CTT20 and H12.2 cells treated with Dox (1 µg/ml, 4 days) were either labeled with EdU for 10 mM alone or further treated with HU (2 mM) for 3 hr and then cells were processed for iPOND assay. Proteins bound to streptavidin beads were analyzed in Western blot. (F) Nuclear dysmorphia increased after NuMA knockdown in H12.2 cells. CTT20 and H12.2 were grown on glass coverslips, treated with Dox (1 µg/ml) for 4 days, fixed with paraformaldehyde, and stained with Lamin B1 to reveal nuclear envelope, and counterstained with DAPI. Abnormal nuclei were defined as nuclei exhibiting clear deviations from smooth, continuous nuclear contours, including blebs (membrane protrusions), invaginations (indentations of the nuclear envelope, INV), or the presence of micronuclei (MN). CTT20 was the control. Scale bars: 5 µm. (G) Percent of nuclear morphology changes in CTT20 cells and in NuMA-depleted H12.2 cells. A total of 288 CTT20 nuclei and 351 H12.2 nuclei were analyzed. Percent of each feature of nuclear abnormality was calculated by dividing the number of nuclei exhibiting a given abnormal feature by the total number of nuclei in each microscopy image. Scoring was performed in a blinded manner.

Interestingly, NuMA depletion did not result in CHK1 S345 phosphorylation (Fig. 4A), and had no effect on CHK1 phosphorylation induced by HU treatment (Fig. 4A). Since ATR/CHK1 is activated by RPA binding to ssDNA, we then measured RPA association to forks in NuMA-deficient cells. Our iPOND results showed that RPA binding to replication forks was largely unchanged in NuMA depleted cells compared to the control cells (Fig. 4E). Even under HU-induced replication stress, RPA32 binding to forks increased in both control and NuMA-depleted cells, as expected; however, NuMA depletion did not lead to any additional increase in RPA32 binding (Fig. 4E). Together, these results indicate that NuMA deficiency does not increase ssDNA at forks or activate the ATR-mediated DNA damage response pathway, despite its significant impact on replication fork progression.

While examining NuMA deficient cells under microscope, we noticed striking nuclear morphology changes and micronuclei accumulation upon NuMA depletion. In contrast to the largely spherically-shaped nuclei in control cells, NuMA-deficient cells exhibited pronounced nuclear dysmorphia including blebs, invagination, and micronuclei (Figs. 4F, 4G), consistent with previous reports showing that loss of NuMA leads to nuclear deformation (Rajeevan, Keshri et al. 2020, van Toorn, Gooch et al. 2023). These structural abnormalities have been linked to defects in nuclear mechanics that can promote DNA damage associated with replication forks (Shah, Hobson et al. 2021). Thus, the genomic instability observed upon NuMA depletion may, at least in part, arise from compromised nuclear integrity.

### NuMA regulates key DNA replication proteins binding at replication forks

The presence of NuMA at replication forks and its impact on regulating DNA replication speed prompted us to investigate whether NuMA could exert any effect on replication machinery. Since NuMA is a structural protein important for maintaining nuclear integrity, we speculated that NuMA dysfunction could destabilize replication protein binding to replication forks. Using iPOND, we measured the localization of proteins related to DNA replication during normal DNA synthesis in NuMA depleted cells. We found that the association of key replication proteins PCNA and DNA Polymerase δ (POLD3) to newly synthesized DNA significantly decreased in NuMA-depleted cells compared to the control cells (Fig. 5A). Such decrease was not due to protein expression change because NuMA depletion did not reduce POLD3 and PCNA protein levels (Fig. 5B). We next tested whether this effect was cell line-specific using an alternative cell line, U2OS. Since iPOND requires large number of cells (2X 10^8^ to 10^9^ cells per sample (Dungrawala and Cortez 2015), performing iPOND in multiple cell lines with siRNA knowdown would be technically challenging and costly. Therefore, we used the SIRF assay to examine the association of replisome proteins with replicating DNA in U2OS cells. Consistent with our iPOND result, NuMA silencing reduced POLD3 binding to forks in the SIRF assay (Figs. 5C, 5D). Again, POLD3 expression was largely unchanged after NuMA depletion in U2OS (Fig. 5E). Since SIRF signals are influenced directly by EdU incorporation efficiency, we performed EdU-SIRF, and observed no significant change in EdU incorporation efficiency in NuMA depleted cells (Suppl Fig. S4). Thus, these results support that NuMA deficiency reduces POLD3 binding to forks. We then examined other key replisome proteins, including POLA1 (POLα catalytic subunit), POLE3 (POLε subunit 3), and MCMs (MCM4 and MCM5). The SIRF signals of these proteins all significantly decreased after NuMA knockdown (Figs. 5F, 5G). Protein levels of POLE3, MCM4, and MCM5 remained largely unchanged upon NuMA depletion (Fig. 5E). However, NuMA depletion slightly decreased POLA1 expression (Fig. 5E). Together, these data suggest that NuMA likely plays an important role in stabilizing or retaining multiple replisome proteins including POLδ, POLε, and MCMs onto replicating DNA. The reduced association of key replisome proteins to replicating DNA may explain the slowed DNA replication speed and increased origin firing observed in NuMA deficient cells (Fig. 3).

**Fig. 5.**
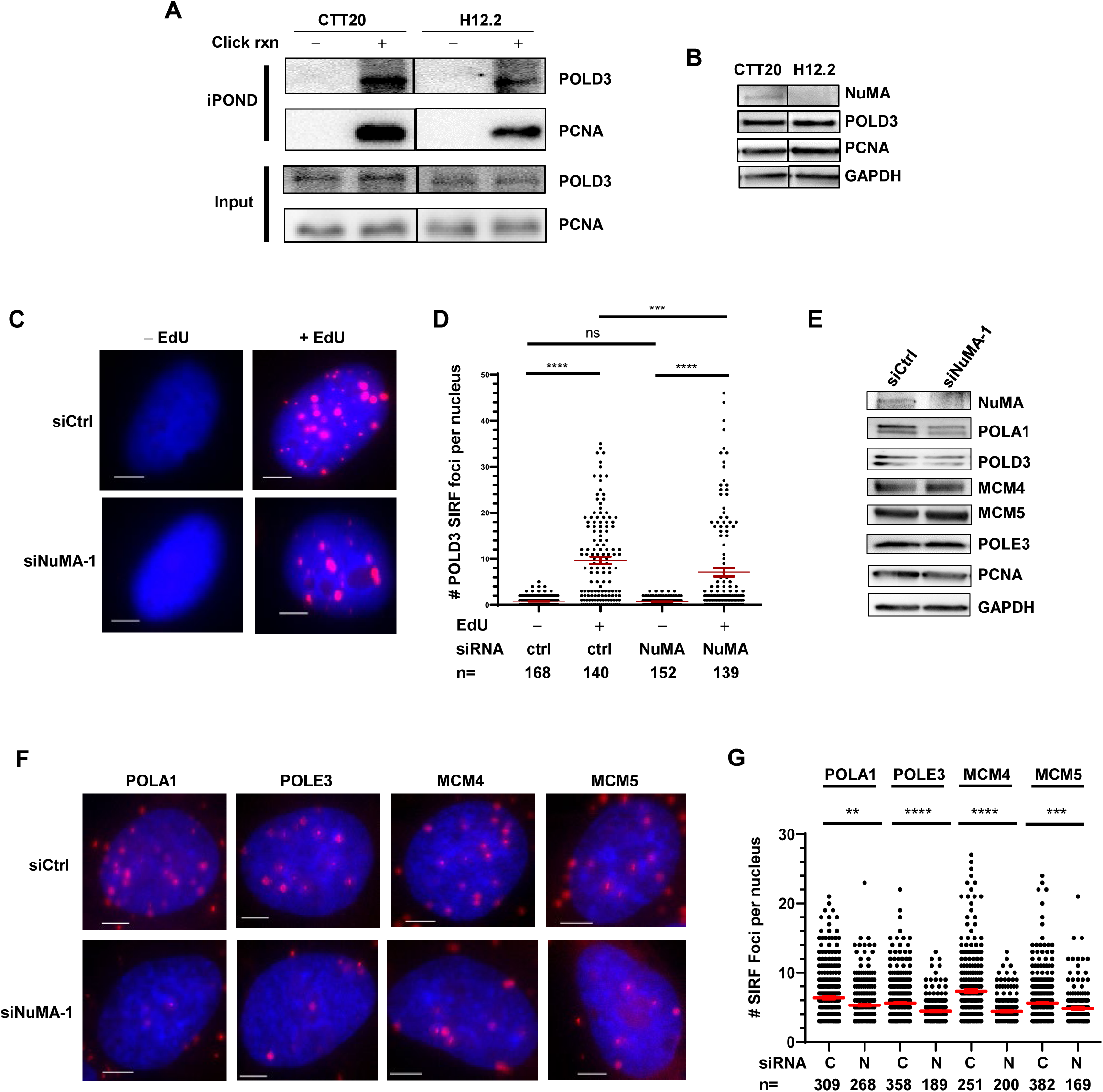
NuMA deficiency impairs replication protein association with forks. (A) iPOND analysis of replisome proteins PCNA and POLD3 in CTT20 and H12.2 cells. Cells were treated as in Fig. 2C. (B) Total protein levels of POLD3 and PCNA were unchanged after NuMA depletion. Cells were treated as in A and whole cell lysates were used for Western blot. (C) Representative SIRF images of POLD3 in U2OS cells upon NuMA depletion with siRNA. U2OS cells were transfected with control siRNA or siRNA targeting NuMA. After 48 hours, cells were processed for SIRF analysis. Scale bars: 5 µm. (D) Quantification of POLD3 SIRF foci upon NuMA depletion. Three independent experiments were performed and result from one experiment is shown. The number of SIRF foci per nucleus were automatically counted with the ZEN software. n: the number of nuclei analyzed. *P* values were calculated using Mann-Whitney test. **** *P*< 0.0001; *** *P*<0.001. Red lines indicate the mean values. Error bars: SEM. (E) Total levels of replication proteins after NuMA knockdown in U2OS cells. NuMA was depleted with siRNA and whole cell lysates were used for Western blot. (D) Representative SIRF images of POLA1, POLE3, MCM4 and MCM5 in NuMA-depleted U2OS cells. Scale bars: 5 µm. (E) Quantification of POLA1, POLE3, MCM4 and MCM5 SIRF foci upon NuMA depletion. C: control siRNA. Two independent experiments were performed and result from one experiment is shown. The number of SIRF foci per nucleus were automatically counted with the ZEN software. N: NuMA siRNA. n: the number of nuclei analyzed. *P* values were calculated using one-way ANOVA. **** *P*< 0.0001; *** *P*<0.001; ** *P*<0.01. Red lines indicate the mean values. Error bars: SEM.

## DISCUSSION

In this study we discovered a novel function of NuMA, a protein primarily known for its roles in mitotic spindle organization and nuclear structure maintenance, in maintaining normal DNA replication. Using both iPOND and SIRF assays, we have observed that NuMA localizes at active DNA replication forks (Fig. 2). Our DNA fiber analysis reveals that NuMA plays an important role in maintaining normal DNA progression and suppressing aberrant origin firing (Fig. 3). In the absence of NuMA, replication forks become susceptible to damage (Fig. 4). These observations identify NuMA as a previously unrecognized regulator of the DNA replication and link NuMA’s known structural roles to the dynamic process of genome duplication, and suggest that beyond its established functions in mitosis and nuclear architecture, NuMA is critical for maintaining normal fork progression and genome integrity.

The localization of NuMA to replication forks, together with the reduced fork speed in NuMA knockout cells, indicate an important role of NuMA in supporting ongoing DNA synthesis rather than an indirect consequence of altered mitosis alone. While the molecular mechanism underlying this slowed progression remains unclear, our results suggest that it may be due to deficiency in the stable assembly of the replication machinery and/or the retention of key replication proteins at forks, as demonstrated by the impaired localization of multiple essential DNA replication proteins like DNA polymerases, PCNA, MCMs at the replication forks upon NuMA loss (Fig. 5). Given NuMA’s known role as a structural protein, we speculate that NuMA may function as a scaffold or organizer that couples higher-order nuclear architecture to the local assembly or stability of replisome components, ensuring efficient and timely DNA synthesis. Disruption of this scaffolding or organization function would result in frequent replisome disassembly or pausing, leading to the slower replication rate and ultimately compromising the fitness of proliferating cells. An alternative explanation for the observed phenotypes is that depletion of NuMA broadly perturbs nuclear architecture, thereby indirectly affecting DNA replication by altering the physical environment in which replication occurs. Given that NuMA is a major structural component of the nuclear matrix, such global effects are plausible and could, in principle, influence multiple chromatin-based processes. However, several observations argue against such a purely nonspecific structural model. First, NuMA loss leads to a distinct combination of increased origin firing and reduced fork progression, rather than a uniform suppression of replication activity, suggesting dysregulation of replication control mechanisms. Second, the preferential activation of ATM signaling over ATR is not typical of canonical replication stress induced by generalized impediments to fork movement, implying a more specific perturbation. We therefore favor a model in which NuMA contributes to the spatial or organizational regulation of DNA replication, although we cannot exclude that part of the phenotype arises from broader changes in the physical environment of the nucleus. Elucidating the precise mechanism by which NuMA influences replication dynamics will be an important direction for future studies.

The observation that NuMA deficiency simultaneously slows replication fork progression and increases origin firing is consistent with the established model that increased origin firing is a compensatory response to fork slowing (Rhind 2006, Zhong, Nellimoottil et al. 2013, Arbona, Goldar et al. 2018). Although it remains to be elucidated how NuMA controls origin firing and replication fork progression, we think it likely helps maintain an optimal balance between the number of active forks and their progression rate by organizing replication factories in defined nuclear microenvironments. Previous studies have demonstrated that proteins that maintain nuclear envelope and internal structure, such as nuclear lamin A/C and associated factors, are in close proximity to replication forks and required for maintaining proper spatial organization of chromatin, which in turn regulates the timing of replication initiation (Dechat, Pfleghaar et al. 2008, Shumaker, Solimando et al. 2008, Lovejoy, Nagarajan et al. 2023, Zhang, Zhao et al. 2025). Notably, NuMA has been reported to associate with the lamin scaffold, and studies have described a partial overlap or colocalization with lamin A/C at the nuclear periphery (Barboro, D’Arrigo et al. 2002, Yamauchi, Kiriyama et al. 2008). It is possible that loss of NuMA may affect lamin A/C-mediated function and cause global structural defects that make chromatin less permissible to rapid replication fork movement, or it could destabilize the physical tethering required to keep the replisome assembled. These findings therefore support a view in which the nuclear structural function of NuMA coordinates origin firing and fork speed to maintain robust DNA replication, analogous to how lamins and other nuclear architectural proteins regulate replication timing and fork stability through modulating chromatin organization. Further studies are needed to delineate how the physical scaffolding provided by NuMA integrates with the biochemical steps of DNA synthesis.

We have observed that loss of NuMA induced increased γH2AX associated with the activation of the ATM/CHK2 DNA damage response pathway, but not the ATR/CHK1 pathway (Figs. 4A-C). Consistent with no ATR/CHK1 activation, our iPOND analysis revealed no increase in RPA binding to forks upon NuMA loss (Fig. 4E), indicating no increase in ssDNA formation at replication forks. This pattern is inconsistent with classical fork uncoupling, in which helicase-polymerase decoupling generates the ssDNA intermediates required for ATR activation. Interestingly, our results show that there is no increase of fork collapse after NuMA depletion (Supple Fig. S3). One possible explanation is that the increased density of replication forks arising from elevated origin firing creates topological stress, potentially leading to low-level, stochastic double-strand break formation at a subset of forks. Given that even a relatively small number of breaks can be sufficient to produce detectable γH2AX and pCHK2 signals, this model could account for the observed ATM pathway activation in the absence of overt fork collapse. Taken together, these findings raise the possibility that NuMA plays a role in coordinating replisome stability and origin firing, and that its loss may shift the replication program toward a break-prone, ATM-activating form of replication stress.

In summary, our findings establish NuMA as a novel component that regulates DNA replication machinery and safeguards genome integrity by promoting efficient replication and preventing the formation of DSBs that trigger the ATM-Chk2 pathway. Future investigations are needed to determine whether NuMA directly interacts with replisome components, and specifically testing the effect of NuMA loss on higher-order chromatin organization within the nucleus to fully validate the proposed link between its structural role and replication maintenance.

## MATERIALS and METHODS

### Cell culture and reagents

HeLa (ATCC #CCL-20TM), U2OS (ATCC #HTB-96) and BJ/hTERT (ATCC #CRL-3627) cells were obtained from American Type Culture Collection. These cells were grown in Dulbecco’s Modified Eagles Medium (DMEM, GE Healthcare Life Sciences, Logan, UT, USA) supplemented with 10% fetal bovine serum (FBS) (Atlanta Biologicals) at 37°C with 5% CO2. NuMA inducible knockout cell lines H11.2 and H12.2 along with the control HeLa cell line CTT20 were gifts from Dr. Cheeseman (McKinley & Cheeseman, 2017), and were cultured in DMEM/F-12 50/50 medium (Corning, 10-092-CV) supplemented with 10% Tet-approved FBS (Biowest, S162TA) and appropriate antibiotics.

The following reagents were used: hydroxyurea (Thermo Fisher Scientific, A10831), EdU (Lumiprobe, 10540), CldU (MP Biomedicals, 105478), IdU (Millipore Sigma, 54-42-2), Biotin-azide (Jena Bioscience, CLK-1265-5), Supersignal West Femto (Thermo Fisher Scientific, Cat# 34095), Supersignal West Pico (Thermo Fisher Scientific, Cat# 34078), Protease inhibitor tablets (Thermo Scientific, A32963), PhosStop Easypack (Roche, 4906845001).

### Immunofluorescent microscopy (IF)

IF was carried out as published previously (Chastain, Zhou et al. 2016). Briefly, cells were grown on the chamber slides overnight, fixed in 4% paraformaldehyde (PFA) for 15 min, washed with PBS three times 5 min each, followed by permeabilization with 0.15% Triton X-100 for 15 min. Cells were then washed with 1X PBS three times 5 min each, blocked with 10% BSA at 37 °C for 1 h in a humidified chamber, and then incubated overnight at 4 °C with primary antibodies, followed by washing with PBS three times 5 min each. Cells were then incubated with secondary antibodies at room temperature for 1 h, followed by sequential washing with either PBS three times 5 min each (for NuMA staining), or PBS once 5 min, Buffer A (0.01 M Tris, 0.15 M NaCl and 0.05% Tween-20, pH7.4) once 10 min, Buffer B (0.2 M Tris and 0.1 M NaCl, pH7.4) once 10 min, 1:100 diluted Buffer B once for 1 min (for γH2AX and Lamin B1 staining). Slides were air dried, and mounting media containing DAPI (Vector Laboratories) were applied. Zeiss AxioImager M2 epifluorescence microscope was used to take the images at 40× or 100×. The following primary antibodies were used: anti-NuMA (1:250, Santa cruz, sc-522268), anti-H2AX pSer139 (1:2000, Active Motif, 39117), anti-Lamin B1 (12987-1-AP, Proteintech, 1:4000). Secondary antibodies were: goat anti-Mouse Alexa Fluor 488 (1:1000, ThermoFisher, A11029), goat anti-rabbit Alexa Fluor 550 (1:1000, ThermoFisher, 84541).

### Transfection of cells with siRNA

Two siRNA sequences targeting NuMA were purchased from Horizon Discovery (#1 Sense: 5’ CCUUGAAGAGAAGAACGAAAUdTdT 3’ and antisense: 5’ AUUUCGUUCUUCUCUUCAAGGdTdT 3’, #2 Sense: 5’ CCACAUCUGAAGACCUGCUAUdTdT 3’ and antisense 5’ AUAGCAGGUCUUCAGAUGUGGdTdT 3’). Negative control siRNA was from Qiagen (Cat# 1022076). Cells were transfected with siRNA oligos with RNAiMax reagents (ThermoFisher, Cat# 13778030) at a final concentration of 80 pmol per well in 6-well plate according to the manufacturer’s protocol. Cells were collected 48 or 72 hours after transfection for fiber assay, SIRF assay or protein analysis.

### Western blot analysis

Western blotting was performed as described (Mahmood and Yang 2012). Briefly, Cells were lysed in pre-chilled lysis buffer (1% SDS, 50 mM Tris-HCl, pH 8.0) supplemented with protease and phosphatase inhibitors, either directly on the dish on ice or after trypsinization followed by lysis of cell pellets on ice. Lysates were sonicated on ice until no longer viscous.

Protein concentrations were determined, and equal amounts of protein were mixed with 2× Laemmli buffer, heated at 95°C for 5-10 min, and resolved by SDS-PAGE. Following transfer to PVDF membranes, membranes were blocked with 5% non-fat dry milk in 1× PBST (PBS with 0.05% Tween-20) for 1 h and then incubated with primary antibodies at 4°C overnight. Membranes were washed with 1× PBST three times 5 min each, incubated with secondary antibodies at room temperature for 1 h, washed with 1× PBST three times 5 min each. The following primary antibodies were used: anti-CHK1 (1:2000, Santa cruz, sc-8408), anti-p-CHK1 Ser345 (1:1000, Cell Signaling, 2348), anti-p-CHK2 T68 (1:1000, Cell Signaling, 2661T), anti-CHK2 (1:1000, Cell Signaling, 2662), anti-GAPDH (1:1000, Cell Signaling, 5174), anti-H2AX pSer139 (1:2000, Active Motif, 39117), anti-H2AX (1:2000, ProteinTech, 10856-1-AP), anti-NuMA (1:1000, ProteinTech, 16607-1-AP), anti-PCNA (1:1000, Santa Cruz, sc-56), anti-POLD3 (1:2000, ProteinTech, 21935-1-AP), anti-RPA32 (1:2,000, Bethyl, A300-244A). Secondary antibodies used are: HRP goat anti-rabbit (WB 1:10,0000, Vector Laboratory, PI-1000), HRP goat anti-mouse (WB 1:5000, BD Pharmingen, 554002). Chemiluminescent signals were obtained with SuperSignal Femto (ThermoFisher) and were acquired using iBright^TM^ FL 1500 Imaging system (ThermoFisher).

### Cell viability assay

CTT20, H11.2 or H12.2 cells were grown in the absence or presence of Dox (1 µg/ml) for 3 days, and then 100 cells were seeded in each well of 6-well plates in triplicates and continued to grow with or without Dox (1µg/ml) in the media for 24 hours. Media were then removed, and 2 ml fresh media without Dox were added to each well and cells were continued to grow for 10 days. Colonies were stained with a 0.2% crystal violet in 50% methanol. Images were taken using iBright^TM^ FL 1500 Imaging system (ThermoFisher).

### Cell cycle analysis

CTT20, or H12.2 cells were grown in the absence or presence of Dox (1 µg/ml) for 4 days, collected by trypsinization, and washed with ice cold PBS. Cells were then resuspended in 50 µl of PBS, followed by the addition of 1 ml of ice-cold ethanol, and stored at 4°C. Prior to FACS analysis, cells were washed with PBS and resuspended in 0.5 ml propidium iodide (PI) staining solution (50 µg/ml PI and 100 µg/ml RNaseA in PBS, 0.1% Triton X-100), incubated at 37°C for 30 min in the dark, followed by addition of 0.5 ml PBS and filtration through a 40-70 µm mesh. Cells were vortexed immediately before acquisition to minimize cell clumping, and flow cytometric data were acquired on a Cytek Aurora spectral flow cytometer using SpectroFlo software (Cytek Biosciences, Fremont, CA, USA). For cell cycle analysis, PI fluorescence was acquired using the YG4 detector/channel according to the instrument configuration. Single cells were gated by FSC-A/FSC-H doublet discrimination before quantification of G0/G1, S, and G2/M populations. Cell cycle distribution was modeled using the Watson Pragmatic algorithm in FlowJo software (BD Life Sciences/FlowJo, LLC, Ashland, OR, USA).

### SIRF assay

SIRF assays were carried out as described previously (Roy, Luzwick et al. 2018). Exponentially growing cells were seeded on the chamber slides and labeled with 125 μM EdU for 8 min. Cells were then washed with PBS once and pre-permeabilized with 0.25% Triton X-100 for 2 min only for the POLD3 antibody, followed by 4 % paraformaldehyde (PFA) fixation for 15 min at room temperature. Chamber dividers were then removed, and cells were washed in PBS three times at room temperature, followed by treatment with 0.25% Triton X-100 for 15 min at room temperature. Click reaction (2 mM copper sulfate, 10 µM biotin azide, and 100 mM sodium ascorbate) was then performed in a humidified chamber for 1 h at 37 °C, followed by washing with PBS three times. In a humidified chamber, cells were blocked with the blocking buffer (10% BSA, 0.1% Triton X-100) at 37 °C for 1 h followed by incubation with primary antibodies at 4 °C overnight. The following antibodies were used in SIRF assays: rabbit anti-biotin (SIRF 1:200, Cell Signaling, 5597), mouse anti-biotin (SIRF 1:200, Sigma, #SAB4200680), anti-POLD3 (1:200, ProteinTech, 21935-1-AP), anti-MCM5 (1:500, Proteintech, 11703-1-AP), anti-MCM4 (1:200, ProteinTech, 13043-1-AP), anti-POLA1 (1:100, Bethyl, A302-851), anti-POLE3 (1:200, ProteinTech, 15278-1-AP). PLA reactions were then performed using Duolink in situ detection reagents red (DUO92008-100RXN, Sigma-Aldrich) following the manufacturer’s protocol. The slides were air dried and mounted with mounting media containing DAPI (Vector Laboratories, H-1200). Images were acquired using the Zeiss AxioImager M2 epifluorescence microscope at 40X magnification. Images were analyzed using the ZEN software (Carl Zeiss AG) and graphs were plotted with GraphPad Prism. The images were analyzed independently by 2 individuals in the lab to avoid human bias. To ensure reproducibility, at least two independent experiments were performed.

### iPOND assay

iPOND experiments were performed as described previously by Dungrawala and Cortez (Dungrawala and Cortez 2015). Briefly, CTT20 and H12.2 cells were grown in the Tet-approved FBS/DMEM/F-12 media with Dox (1 µg/ml) for 4 days. Cells around 80-90% confluency in 150mm dish were labeled with EdU (10 µM) for 10 min, then either treated with HU (2 mM) for 3 hours or untreated, fixed with 10 ml of 1% formaldehyde for 20 min at room temperature. After neutralization by addition of 1 ml of 1.25M glycine, cells were scraped from plates and washed with 1X PBS for 3 times at 4°C. After permeabilization of cells, cell nuclei were subjected to click reaction with 10 mM sodium ascorbate, 2 mM CuSO4, plus or minus 10 µM biotin picolyl azide (Click Chemistry Tools, Cat # 1167-5) in PBS for 1-2 hours with constant rotation at RT. Cells were then lysed in lysis buffer (1% SDS/50 mM Tris-HCl, pH8.0) in the presence of protein inhibitor cocktail and sonicated 6-10 times with a power of 10 Watts, 15 sec pulse. 1% of supernatants were taken out as INPUT, and the remaining supernatant was incubated with High-capacity Streptavidin Agarose Resin (ThermoFisher Scientific, 20357) overnight in the dark. The beads were washed once with cold lysis buffer, once with 1 M NaCl, 2 times with lysis buffer and then incubated with 2X SDS Laemmli buffer with 0.2M DTT and heated at 90°C for 25 min. The eluted proteins bound to captured DNA were separated in SDS-PAGE gel and transferred to PVDF membrane for detection of different proteins with specific antibodies.

### DNA fiber assay

DNA fiber assay was carried out using the previously published protocol (Nieminuszczy, Schwab et al. 2016). For normal DNA replication, 70% confluent cells were pulse-labeled with the thymidine analogs 50 µM 5-chloro-2’-deoxyuridine (CldU) (MP Biomedicals 105478) for 20 min. CldU was then removed by washing twice with pre-warmed PBS and nascent DNA was further labeled with 250 µM 5-iodo-2’-deoxyuridine (IdU) (Millipore Sigma 54-42-2) for 20 min. Cells were washed with PBS to remove CldU and IdU and collected by trypsinization. After harvesting, cells were washed once with PBS and resuspended at a density of ∼1000 cells/µl in PBS. Cell suspension (2 µl) were dropped onto a glass slide and allowed to become half-dry in 5 min. Then 12 µl of lysis buffer (200 mM Tris-HCl pH 7.5, 50 mM EDTA, 0.5% SDS) were added to cells on glass slides and stirred cells in circular motion. Slides were then put in a humidified chamber for 2 min for efficient lysis. Slides were then inclined to 15° to allow genomic DNA spread. Spread DNA fibers were air dried for about 30 min and then fixed in methanol: acidic acid (3:1) for 10 min. Slides were then immersed into 2.5 M HCl for 100 min to denature DNA and then washed with 1× PBS three times. Slides were then blocked with 5% BSA for 30 min, immunostained with rat anti-BrdU 1:500 (Abcam, ab6326, cross-reacts with CldU) and mouse anti-BrdU 1:50 (BD Biosciences, 347580, cross-reacts with IdU) primary antibodies for 1 h in a humid chamber at 37 °C. After washing with PBS three times, slides were incubated with secondary antibodies anti-rat Alexa Fluor 488 1:500 (ThermoFisher Scientific, A11006) and anti-mouse Alexa Fluor 568 1:500 (ThermoFisher Scientific, A11031) at 37 °C for 1 h. After washing with PBS and air dry at room temperature in dark, slides were mounted in the mounting medium without DAPI (Vector Laboratories, H-1000) using coverslips. Images were acquired using the Zeiss AxioImager M2 epifluorescence microscope at 40× magnification. DNA tract lengths of CldU (green) and IdU (red) were measured and analyzed by the ZEN software (Zeiss, Carl Zeiss AG). About 100-200 fibers were analyzed for each sample in each experiment and data were analyzed by GraphPad Prism. To obtain fork speed in kb/min, the length of CldU or IdU tracks in kb was divided by the labeling time (20 min). Fiber length values were converted into kb using the conversion factor 1 µm = 2.59 kb (Jackson and Pombo 1998). When determining replication origins, stalled or termination fibers (green and green-red-green) were excluded from counting. Origins that fired during the 1st analog labeling (CldU) were identified as red-green-red events. Origins that fired during the 2nd analog labeling (IdU) were identified as isolated red events. The percentage of the 1st label origins (red-green-red) was calculated using the following formular: % of the 1st label origins = (all red-green-red tracks/all green (CldU)-labeled tracks) x 100. The percentage of the combined 1st and 2nd label (red-green-red and red only) origins was calculated using the following formula: % of (1st + 2nd label origins) = (all red-green-red tracks + red only tracks)/(total green (CldU)-labeled tracks + red only tracks) x 100. Inter-origin distance was calculated as the distance between the midpoints of two neighboring green tracks from red (IdU) - green (CldU) - red (IdU) - green (CldU) - red (IdU) labeled fibers. Percentage of green-only fiber was calculated by dividing the number of green-only tracks by the total number of green tracks in the image.

### Statistical analysis

Graphpad Prism 10 was used for statistical analysis. The data are presented as the mean ± standard error, and the statistical significance of the difference between subjects was evaluated with Mann-Whitney test for comparing the significance between two subjects, one-way ANOVA test for comparing the significance among more than two subjects. Statistical significance is indicated for each graph (ns = not significant, for *P* > 0.05; * for *P* < 0.05; ** for *P* < 0.01; *** for *P* < 0.001; **** for *P* < 0.0001).

## Author contributions

Z-GW, SK and FW performed experiments and data analysis. Z-GW assembled figures. Z-GW and WC wrote the manuscript. WC conceived and supervised the study, and edited the manuscript.

## Funding

This work is supported by National Institute of Health (NIH) R01GM146376 to WC. The funding bodies had no role in study design, data collection, data analysis and interpretation.

## Acknowledgments

We thank Dr. Cheeseman at Whitehead Institute MIT for CTT20, H11.2 and H12.2 cell lines.

## Conflicts of Interest

The authors declare that they have no conflict of interest.

## Data Availability

All data generated or analyzed during this study are included in this article and its supplementary information files.

---

**Fig. S1.**
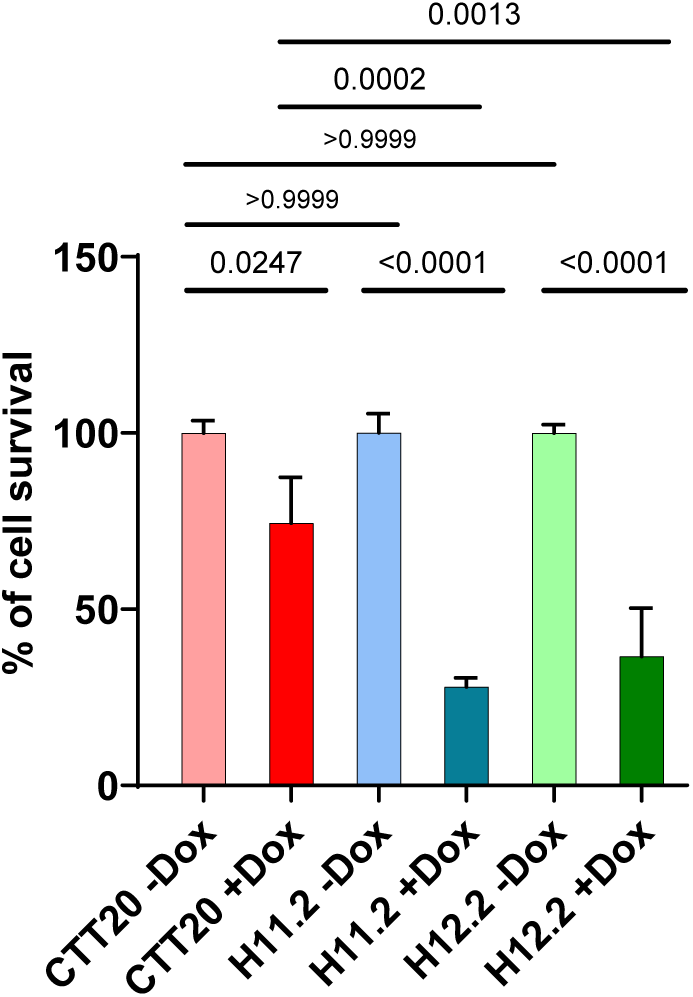
Colony formation analysis of CTT20, H11.2 and H12.2 treated with and without Dox showing that Dox/Cas9 induction modestly reduces cell viability in control cells.

**Fig. S2 (Related to Fig. 4D).**
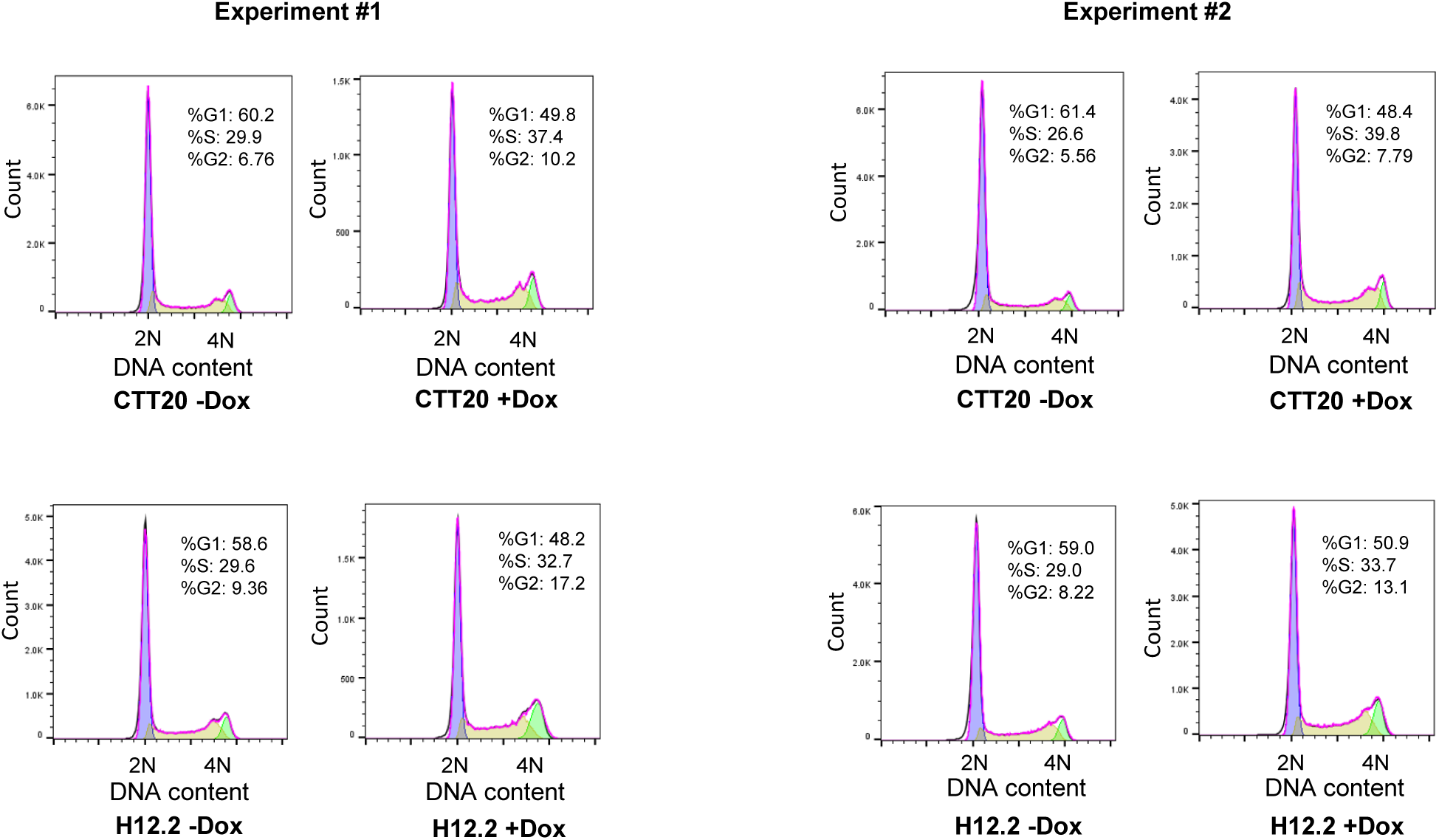
Histogram of cell cycle analysis. CTT20 and H12.2 were grown in the presence/absence of Dox for 4 days and were collected for evaluation of G1, S, and G2 phase by flow cytometry. The flow cytometric images from two experiments (#1 and #2) were shown.

**Fig. S3 (Related to Figure 4).**
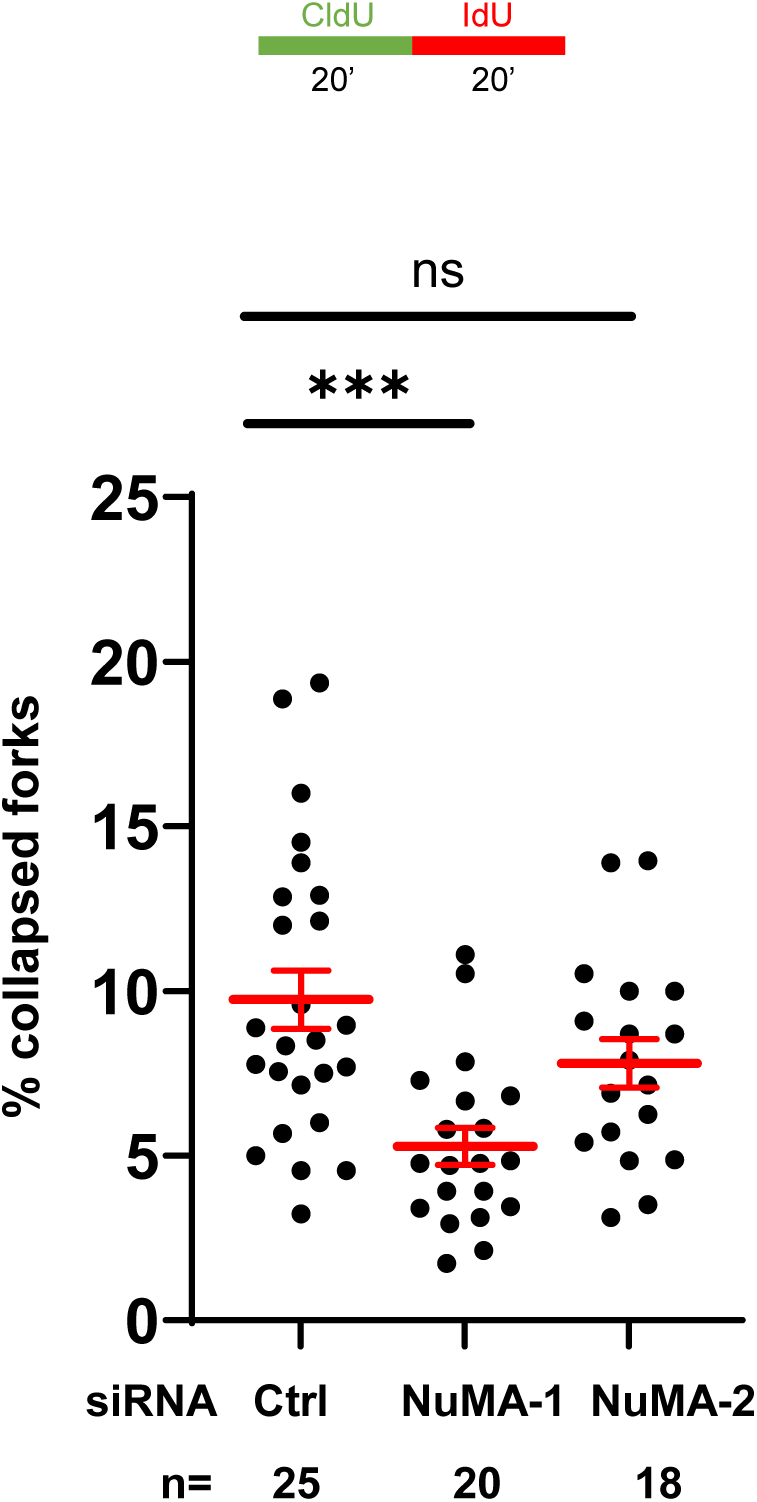
Percent of collapsed forks in NuMA knockdown cells. HeLa cells transfected with control or NuMA siRNA were sequentially labeled with CldU (20 min) and IdU (20 min) for DNA fiber analysis. Percent of stalled forks (CldU-only tracks) was calculated by counting the number of stalled forks in each microscopic field and normalizing to the total number of DNA fibers in the same field. n: the number of individual microscopic images, with each image containing approximately 50 DNA fiber tracks. Red lines indicate mean values. Error bars: SEM. *P* values were calculated using one-way ANOVA.

**Fig. S4.**
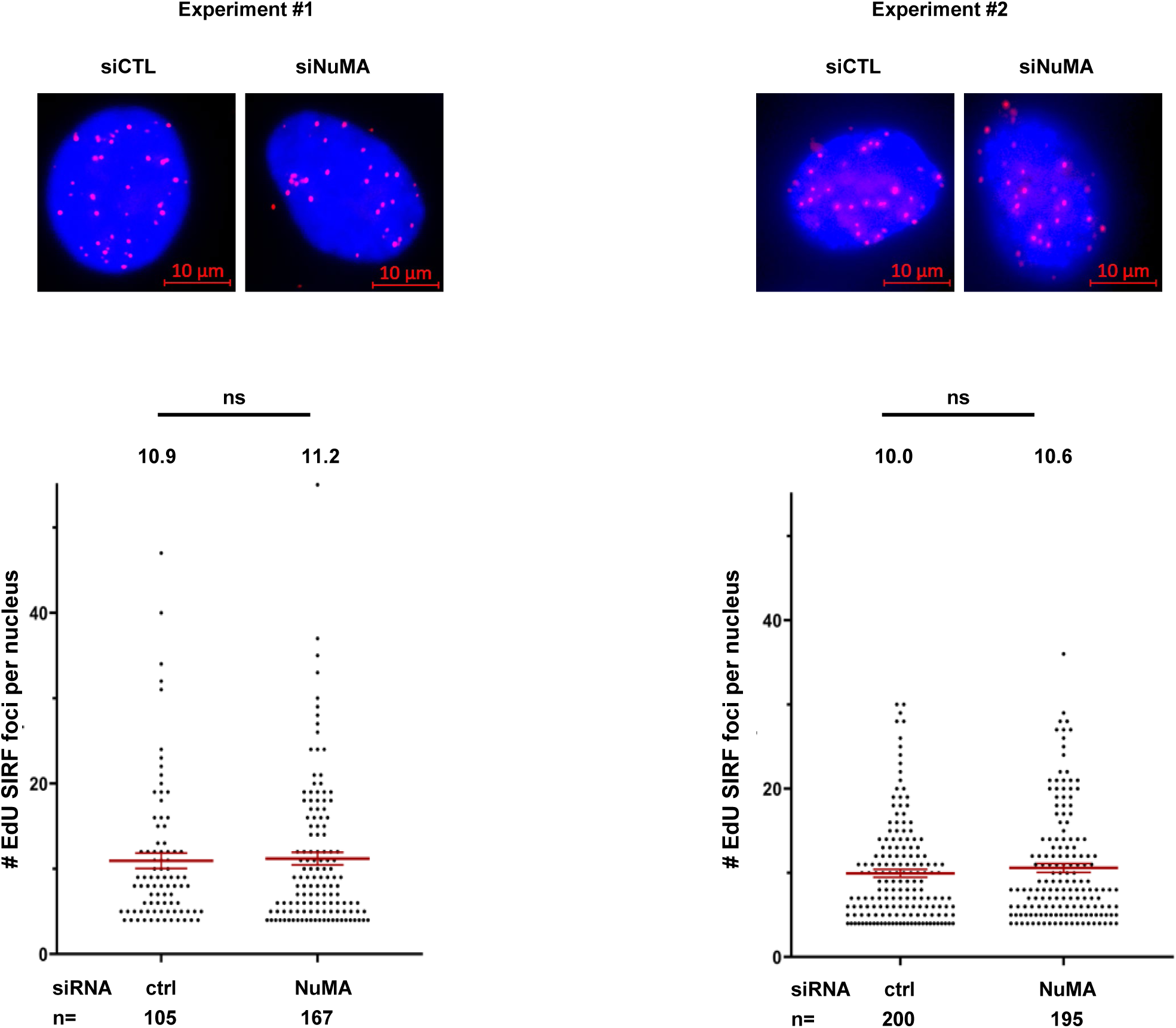
Biological replicates of EdU-SIRF in U2OS cells following NuMA knockdown. U2OS cells were transfected with indicated siRNA to deplete NuMA, pulse labeled with EdU for 8 min, and subjected to SIRF assay as described in Methods. After the click reaction, cells were incubated with two different anti-biotin antibodies (rabbit and mouse), followed by PLA reactions to assess whether NuMA depletion affects EdU incorporation into nascent DNA. Representative EdU-SIRF images from each experiment are shown. The number of EdU-SIRF foci per nucleus was counted. *n* represents the number of nuclei analyzed in each sample. Red lines indicate mean values, which are annotated above the data points. Error bars: SEM. Scale bars: 10 µm.

## Notes

### Competing Interest Statement

The authors have declared no competing interest.

